# Automated assessment of the mouse body-language reveals pervasive behavioral disruption in a two-hit model of psychiatric vulnerability

**DOI:** 10.1101/2025.01.31.635700

**Authors:** Viviana Canicatti, Vanes Cibin, Nicoletta Berardi, Yuri Bozzi, Alessandro Sale, Gabriele Chelini

**Affiliations:** Institute of Neuroscience, National Council of Research, Pisa (IT); Center for Mind/Brain Sciences, University of Trento, Rovereto (IT)

## Abstract

The influence of early-life experiences is widely acknowledged as a crafting tool that sculpts complex behavioral patterns and well-being of living organisms. The use of preclinical models can provide invaluable insight into how a negative environmental push interplays with genetic make-up in shaping psychiatric vulnerability. However, the assessment of psychiatric traits in cross-species studies often relies on the use of surrogate metrics as a proxy for the internal state, limiting the interpretation to context-dependent outcomes. In this work, we exploited a validated computational tool for digitalized ethological screening to identify spontaneous hallmarks of altered behavioral functioning in a dual-hit mouse model of psychiatric vulnerability. To do so, mice carrying heterozygous deletion of the gene coding for Contactin-associated protein-like 2 (*Cntnap2^+/-^*) and their wild-type (WT) littermates were raised with limited bedding and nesting (LBN). These animals were compared to both WT and *Cntnap2^+/-^* mice raised in standard conditions, mapping their spontaneous behavior during freely-moving exploration. Our data show that descriptors of motility state or surrogate anxiety indicators largely failed in detecting subtle diversion from control conditions. By contrast, automated segmentation of the body-language revealed a significant impact of both genotype and early-life experience in shaping the spontaneous behavioral programming. Thus, using unsupervised clustering, we unveiled two alternative neurobehavioral profiles within our dataset. We found that one of the identified profiles largely overlapped with *Cntnap2^+/-^* mice raised with LBN, while the other was equally shared among controls. We conclude that the coincidence of early-life adversity and *Cntnap2* haploinsufficiency drastically reshapes behavioral structure in rodents.

**SIGNIFICANCE STATEMENT:** Enhancing the predictive and face validity of preclinical models in psychiatric research remains a significant challenge due to the inherent heterogeneity and complexity of these conditions. While animal models are crucial for understanding the risk factors involved, replicating the full complexity of these conditions continues to pose difficulties. In this study, we use a tool for digitalized behavioral screening to investigate emotional hallmarks in a double-hit (environmental and genetic) mouse model of vulnerability for psychiatric disorder, glimpsing subliminal behavioral disturbances not captured with traditional assessments. Our findings highlight the effectiveness of novel computational tools in identifying subtle behavioral deviations and support the hypothesis that gene-environment interaction contributes to shape alternative behavioral structure in mice.

## INTRODUCTION

Genetic factors and adversities during early-postnatal life (ELA) are universally recognized as major contributors to psychiatric vulnerability (1). The use of animal model is an ideal way to stratify the relative contribution of genetic and environmental vulnerabilities that shape core behavioral hallmarks of psychiatric disorders (1). However, the majority of currently validated empirical tools to identify psychiatric-like traits in rodents consist in challenging naïve mice with forced-choice tasks, oscillating between safe options and anxiogenic alternatives (3–5). Alternatively, studies can rely on the use of acquired (6, 7) or innate fear memories (8, 9), or induced elevation of stress (10, 11). These approaches constrain the behavioral evaluation to a highly context-dependent scenario, which is not necessarily reflective of the chronically disabling nature of mental illness (12, 13).

To overcome this limitation, in this work we asked whether genetic and environmental vulnerability to multiple psychiatric disorders contribute to the definition of alternative behavioral pattern in rodents, as an indicator of altered neurobehavioral functioning. To do so, we focused on CNTNAP2 heterozygous mutation. Haploinsufficiency of CNTNAP2 in humans was previously associated with multiple psychiatric diagnosis (schizophrenia, bipolar disorder, autism spectrum, major depressive disorder), as well as neurotypical individuals (14–18). By consequence, CNTNAP2 contribution to psychopathology is not fully understood. Applying the logic of the Research Domain Criteria framework (RDoC) (19), we reasoned that *Cntnap2* haploinsufficiency may alter the global neurobehavioral functioning in rodents, rather than impacting isolated domains, thus resulting in a complex vulnerability to environmental hits. Studies show that mice with *Cntnap2* heterozygous mutation are virtually indistinguishable from wild-types (20, 21). Another study, however, proved that, when paired with a prenatal immune activation, *Cntnap2* haploinsufficiency enables the emergence of autistic-like behavior in mice (22). This finding suggests that *Cntnap2* haploinsufficiency interacts with environmental factors to exacerbate behavioral abnormalities in rodents. Here we hypothesized that the exposure to an early environmental stressor may impact animals with C*ntnap2* heterozygous mutation by altering their expression of spontaneous behavior.

To this goal, we took advantage of a widely used tool to induce early-life stress in rodents, the limited bedding and nesting paradigm (LBN) (23–25). Litters composed of both *Cntnap2*^+/+^ and *Cntnap2*^+/-^ raised in LBN (from this point onward defined as *Cntnap2^+/+^LBN* or *Cntnap2^+/-^LBN* respectively) were compared with parallel litters of standard reared *Cntnap2*^+/+^ and *Cntnap2*^+/-^ (defined as *Cntnap2^+/+^*SR or *Cntnap2^+/-^*SR respectively) and evaluated during freely-moving exploration of an open field. Using a computational pipeline for digitalized ethological screening (stimulus-evoked behavioral tracking in 3D for rodents (SEB3R) (2), the mouse body language was partitioned in discrete behavioral modules (BMs) to capture qualitative discrepancies in the expression of naturalistic behavior.

We observed that, while having limited effect on individual BMs, both *Cntnap2* haploinsufficiency and ELA play a role in altering the overall composition of spontaneous behavior. Then, using unsupervised clustering to stratify the study cohort, we discovered that *Cntnap2^+/-^*LBN mice exhibit an alternative and nearly exclusive behavioral structure (i.e. personality) compared to the other groups. Moreover, we noticed that this atypical behavioral expression is characterized by a remarkable inter-individual homogeneity, indicating that the combined effect of *Cntnap2* haploinsufficiency and early-life stress constrains behavioral strategies into highly stereotyped programs.

These findings suggest that *Cntnap2* haploinsufficiency alters the expression of naturalistic behavior in mice, and exacerbates in a drastic reshaping of the spontaneous neurobehavioral functioning when paired with an early-life adverse experience.

## RESULTS

### Traditional metrics did not spot any significant influence of *Cntanp2^+/-^* and LBN on spontaneous behavior

First, we evaluated four experimental groups (*Cntnap2^+/+^* and *Cntnap2^+/-^* LBN with their respective SR controls) using the traditional metrics of a classical open field test (Fig. 1A), looking into the mice preferential choice for the borders and corners (safe option) against the center (anxiogenic option) of the arena. Our data show that none of the group display significant changes in the preference for the corners (Fig. 1B) or the borders of the apparatus (Fig. 1C), indicating a lack of obvious hallmarks of anxiety. Moreover, we found no differences among the four groups in the motility state (Fig. 1D-F). These findings suggest that neither the affective nor the motor domains are impacted during the freely-moving exploration of a neutral environment.

**Figure 1.**
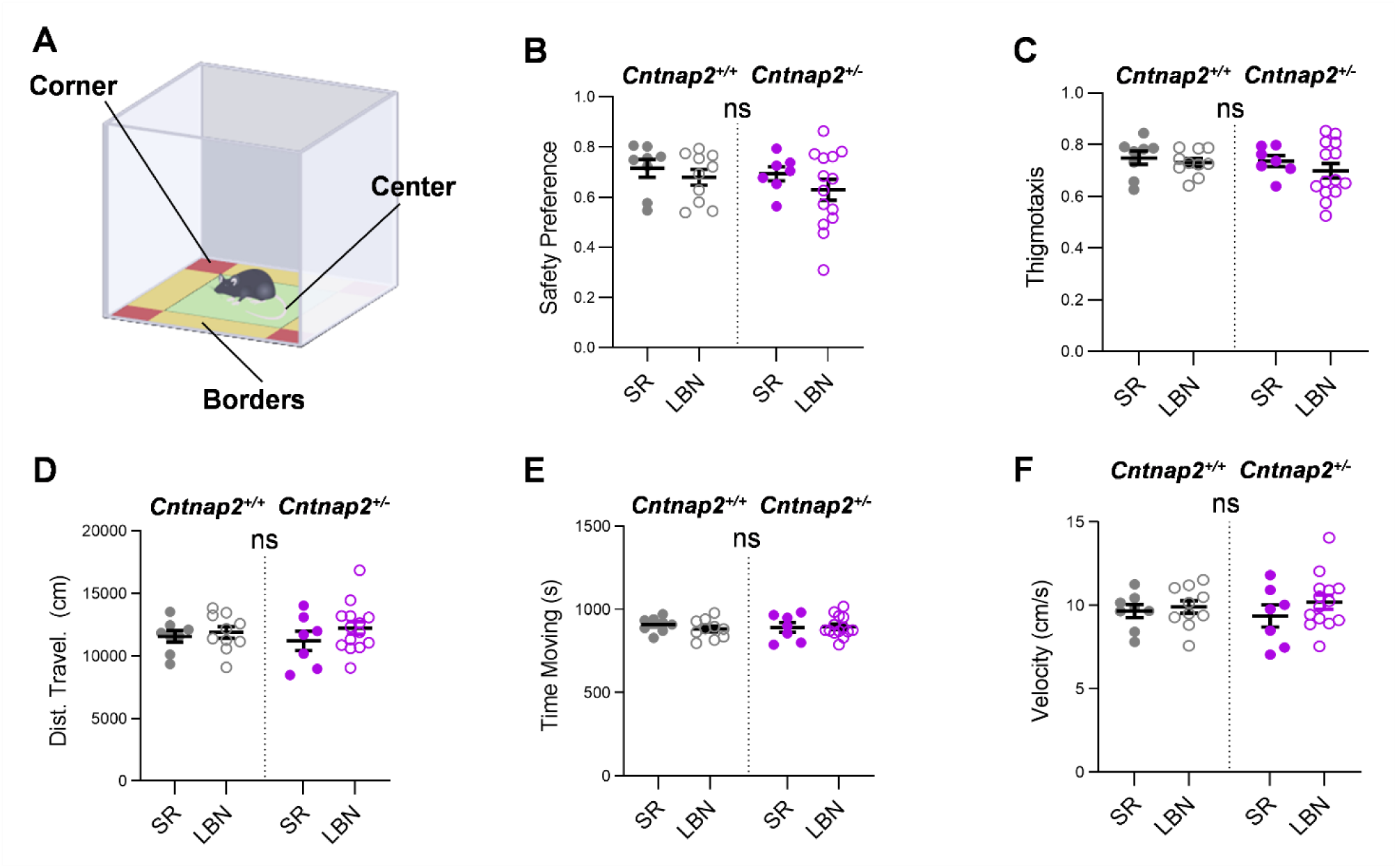
Traditional measures failed in detecting group discrepancies in both anxiety and locomotor activity. **A)** Graphical representation of the Open Field arena. **B)** Neither the rearing condition (ELE) nor genotype has any significant impact on anxiety measurements over the 20-minute of open field (OF) test. **C)** Similarly, no effect of environmental and genetic variables on locomotor activity. Each dot represents an individual. Data are expressed as mean +/- SEM. Statistical analysis was performed by two-way ANOVA (B, C). **ns:** not significant.

### Both *Cntnap2* heterozygous mutation and ELA alter naturalistic behavior in mice

To capture the nuances of behavioral expression we analyzed the same experimental sessions using SEB3R (Fig. 2A, B). With this tool, the mice body-language was deconstructed in a total of nine distinct BMs (Fig. 2C). First, we focused on quantifying difference in individual BMs. Our analysis only detected a limited effect of the variable early-life experience (ELE, p=0.014) and a nearly significant effect of the gene x environment interaction (ELE^X^GEN, p=0.051) on the rearing-associated BM2. The difference was mostly driven by significant reduction of BM2 in *Cntnap2^+/-^*LBN with respect to SR genotype-matched controls (p=0.009). Furthermore, a main effect of the variable genotype (Gen) was found for BM5 (p=0.001), driven by significant reduction of both *Cntnap2^+/-^* groups (SR and LBN) compared to *Cntnap2^+/+^*SR (p=0.02 and p=0.009 respectively, supplementary figure 1E).

**Figure 2.**
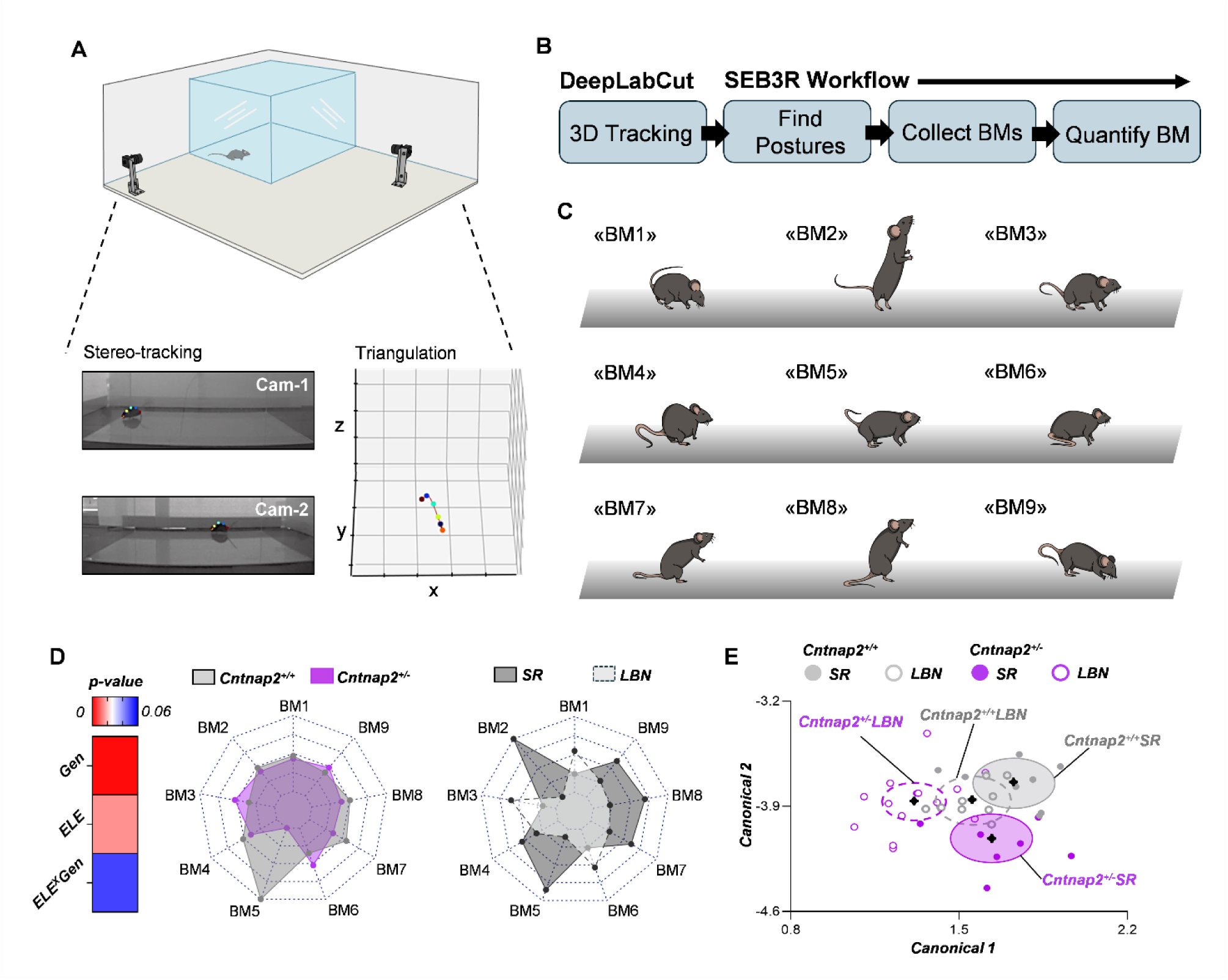
Digitalized ethological screening suggests the role of both *Cntnap2* haploinsufficiency and early-life adversity in shaping the spontaneous behavioral pattern. **A)** Animals were video-recorded in an open field arena by two cameras in stereo-configuration; key points along the mouse spine were tracked in 3D using DeepLabCut. **B)** Workflow of SEB3R pipeline as previously described (2). **C)** SEB3R identified nine Behavioral Modules (BM) during freely moving open-field exploration. **D)** The global expression of BMs is significantly affected by both genotype (Gen, p=0.001) and early-life experience (ELE, p=0.014), while the interaction between the two variables shows a nearly significant effect (ELE^X^Gen, p=0.051). **Left:** heatmap displaying the weighted impact of the variables onto BMs expression. **Right:** radar plots depicting qualitative differences in BMs expression between Genotypes (left) and rearing condition (right). (For visualization purposes, radar plots report data as percentage (%) of change with respect to the wholesome group mean). **E)** Centroid plot displaying the results of 2-way MANOVA when taking into account all the covariates (the ellipses indicate the 95% confidence interval containing each of the groups. Roy’s max root p=0.0092**). Centroids are indicated with the symbol: 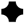

More importantly, when taking into account the overall changes in all BMs expression, we identified a significant effect of both Genotype (p=0.001) and LBN (p=0.01), and a near-to-significance effect of the Gen x LBN interaction (p=0.05) (Fig.2D). This finding was also supported by the visible misalignment of the experimental groups onto the centroid plot obtained by calculating canonical scores (Roy’s largest root p=0.009. Fig.2E). These findings indicate that *Cntnap2* haploinsufficiency plays a significant role in shaping the behavioral structure in rodents and suggest that ELA may differentially impact subjects carrying this mutation.

### Cluster analysis unveils a nearly-exclusive behavioral profile in *Cntnap2^+/-^*LBN

Then we asked whether the discrepancies in BM expression may determine an alternative neurobehavioral functioning in our study cohort. To test this hypothesis, we used BMs expression as classifiers for unsupervised *k-mean* clustering and stratified our dataset into two distinct behavioral profiles (i.e. personalities: P1(*n =* 20) and P2 (*n =* 18). Fig.3A, B). The successful distinction between the two personalities was confirmed using 2-way ANOVA on the Euclidean distances from centroids (Fig. 3C). To elucidate whether our independent variables played a role in determining the alternative behavioral profiles we asked whether any of the identified personalities was preferentially related to either experimental condition. Strikingly, we found that the *Cntnap2^+/-^* LBN group was nearly exclusively characterized by the personality “P2” (84.62% of the total). To the contrary, personality P1 was similarly, and predominantly, represented in the other conditions (87.5% *Cntnap2^+/+^*SR/ 66.67 *Cntnap2^+/+^*LBN / 62.5 *Cntnap2^+/-^*SR). Significant differences in the frequency distribution were confirmed by *χ^2^*test (Fig. 3D, *p*=.004). Specifically, personality P2 displayed significantly reduced rearing-associated BMs (BM7, *p* = 1.07e-05. BM8, *p* = .0004. BM2, *p* = .0002) as well as BM4 (*p* = .005), compared to P1. Conversely, BM3 was significantly reduced in P1 compared to P2 (*p* = .007) (Fig. 3 E, F). These data can be appreciated on a single-subject level (Fig. 3E) or as a group comparison (Fig. 3F).

**Figure 3.**
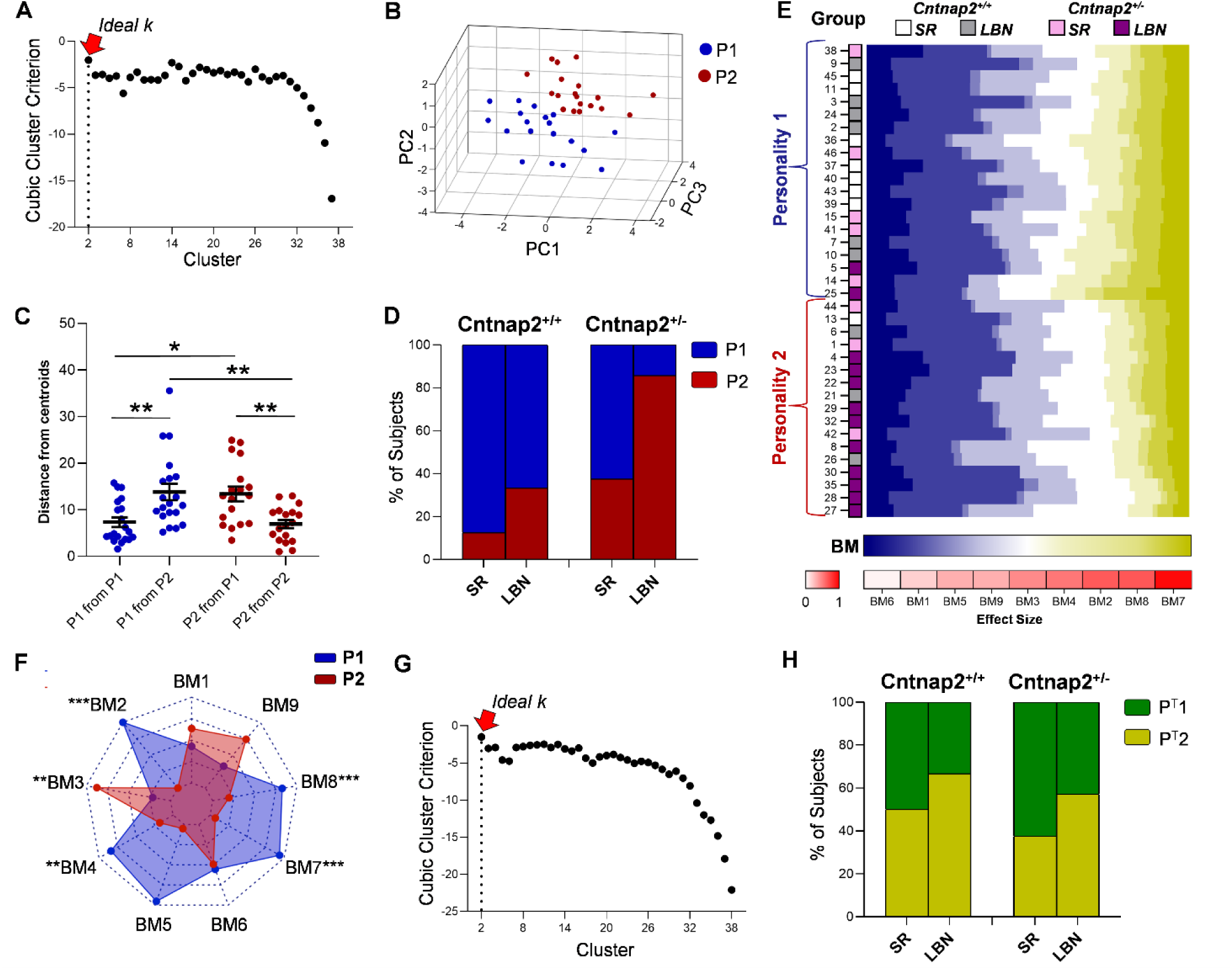
Unsupervised clustering using SEB3R output evidence a distinctive behavioral profile in *Cntnap2*^+/-^ mice raised with LBN. **A)** Graphical depiction of the cubic clustering criterion (CCC) for unsupervised *k-mean* cluster selection using SEB3R output, suggesting the presence of two distinct behavioral profiles [P1 and P2]. **B)** Scatter plot showing the separation of clusters identified using SEB3R output. **C)** Betweenness and Withiness of clusters confirms the efficacy of *k-mean* in separating distinct behavioral profiles (2-way ANOVA, P1-P1 vs P1-P2 p=0.005; P1-P1 vs P2-P1 p=0.012; P2-P1 vs P2-P2 p=0.009; P1-P2 vs P2-P2 p=0.003). **D)** Behavioral profiles are differentially represented among *Cntnap2*^+/-^LBN with respect to any other group (*χ^2^ p= 0.004**).* **E)** Raster plot depicting the density of each BM for each subject. BMs are arranged (left to right) according to their effect size in driving the differences between behavioral profiles (graphically represented in the red gradient map below the plot). Note the drastic shift in BM density between P1 and P2, beginning at BM3. **F)** Polar plot displaying the differences in BMs expression of the two behavioral profiles; BM2,4,7,8 are significantly reduced in P2 compared to P1. BM3 is increased in P2 respect to P1. **G)** Graphical depiction of the CCC method for unsupervised *k-mean* cluster selection using traditional behavioral descriptors suggests the presence of two distinct behavioral profiles [P^T^1 and P^T^2. **H)** Behavioral profiles identified using traditional metrics (P^T^) are equally distributed across experimental groups (*χ^2^ p= 0.61)*. *p<0,05 / **p<0,001 / ***p<0,0001.

Finally, we assessed whether the discrepancies emerging from the analysis of the body language were also reflected in differential expression of classical indicators of stress, anxiety, and locomotor activity. We applied an identical clustering strategy using the parameters obtained from the open field test as classifiers (shown in Fig. 1). Once again, two predominant behavioral profiles emerged (P^T^1 and P^T^2. Fig. 3G), but none of the two showed any privileged association with neither of the experimental groups (*χ^2^ p=0.61, Fig. 3H)*, nor overlap with the previous classification.

These data show that SEB3R analysis captures discrepancies in the naturalistic behavior where classical OF fails to do so, showing that the combined effect of genetic and environmental vulnerability operates by shaping alternative behavioral programs, rather than linearly increasing, or decreasing, the levels of stress and anxiety.

### Subjects expressing personality P2 display a remarkable inter-individual homogeneity

The emerging principles of stratified psychiatry (26) and the RDoC criteria framework (19) suggest that the high heterogeneity observed within diagnostic populations derives from discrepancies in the underlying etiology (19, 26). By translating this principle into our preclinical model, we hypothesized that the personality P2 should present a higher inter-individual homogeneity due to the predominant presence of subjects belonging to the *Cntnap2^+/-^*LBN group. Conversely, P1 may incorporate a scramble collection of behavioral styles, due to discrepancies in the subjects’ genetic and environmental background. To test this hypothesis, we generated two similarity network structures using the weighted pairwise Spearman correlation for each of the two personalities (Fig. 4 A, B). As predicted, we found that the P1 network display a sparse and fragmented structure, with few isolated nodes and polarized interconnections (Fig. 4B-top). The network P2, on the other hand, appeared largely homogenous, with no visible outliers and nodes exhibiting well-organized connections (Figure 4B-bottom). Thus, we extracted and compared the centrality measure of both networks, using them as an indicator of groups homogeneity (Fig. 4 C, D and supplementary Fig. 2), indicating that the expression pattern of BMs is highly stereotyped in P2 and more individualized in P1.

**Figure 4.**
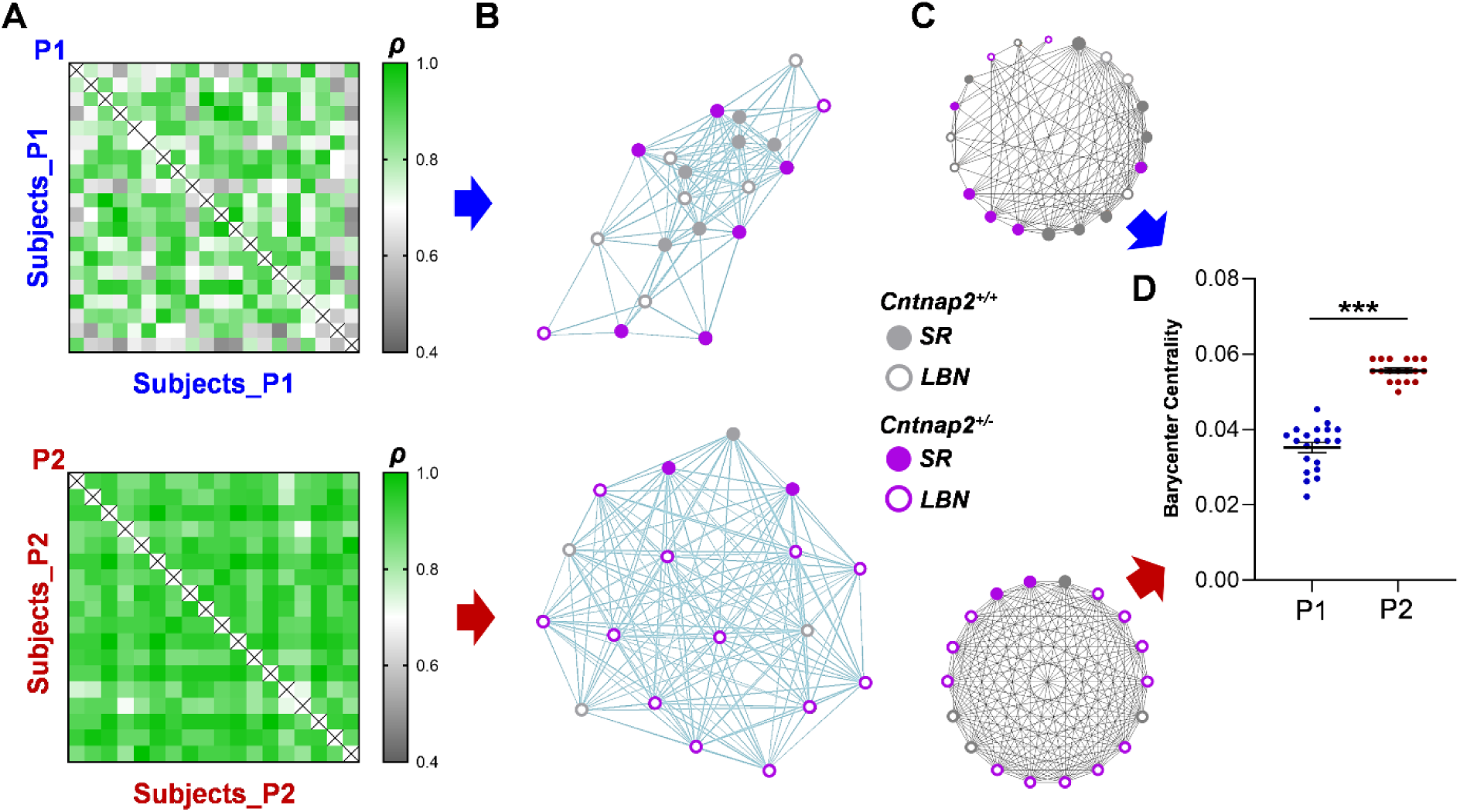
The artificial group P2 exhibits a remarkable intrinsic homogeneity compared to P1. For both groups, the correlation matrix of BM expression **(A)** was transformed into a network **(B)** where each node is a subject and the connecting lines are the pairwise degree of similarity. Then, the centrality measures of the network **(C)** were used as a measure of the group homogeneity. **(D)** Network barycenter centrality is significantly higher in P2 (Mann-Whitney test p>0.0001). ***p<0,0001.

## DISCUSSION

The use of novel computational tools developed by system neuroscience provides the opportunity of integrating traditional behavioral test with advanced methods for detection and analysis of spontaneous behavioral expression in rodents (2, 27–33). In this study we applied a validated computational pipeline for digitalized ethological screening (SEB3R (2)) to identify spontaneous hallmarks of altered behavioral expression in a dual-hit model of psychiatric vulnerability.

We show that mice with heterozygous mutation for *Cntnap2* present atypical body-language expression during freely-moving exploration of a neutral environment. Importantly, our data highlight marginal changes in specific BM, but meaningful discrepancies in the global behavioral structure. Furthermore, the observed changes were so amplified by the impact of LBN, that using a data-driven method for dimensionality reduction led us to pull apart a behavioral profile nearly exclusive for the *Cntnap2^+/-^*LBN group (Fig. 3). These findings suggests that multi-modal vulnerability to mental illness drastically reprograms the neurobehavioral functioning, putatively changing the way a subject interacts with a constantly changing environment.

We also report that, among the two behavioral profiles that emerged, the one more that is more strongly represented among the *Cntnap*2^+/-^LBN (P2) showed a dramatic increase of inter-individual similarity. Reduced cognitive-behavioral flexibility is a common traits in psychiatric disorders, including in conditions associated with *Cntnap2* haploinsufficiency, such as schizophrenia and autisms (12, 34). Based on our findings, we speculate that genetic and environmental vulnerabilities act by limiting the complexity of behavioral strategies in favor of stereotyped programs (Fig. 5). These findings also suggest that stratifying the population on the basis of the spontaneous behavioral expression may implicate similarities in the underlying etiology. This observation corroborates novel interpretative models in biological psychiatric research and speaks to the need of a dimensional neurobehavioral classification (19, 26).

**Figure 5.**
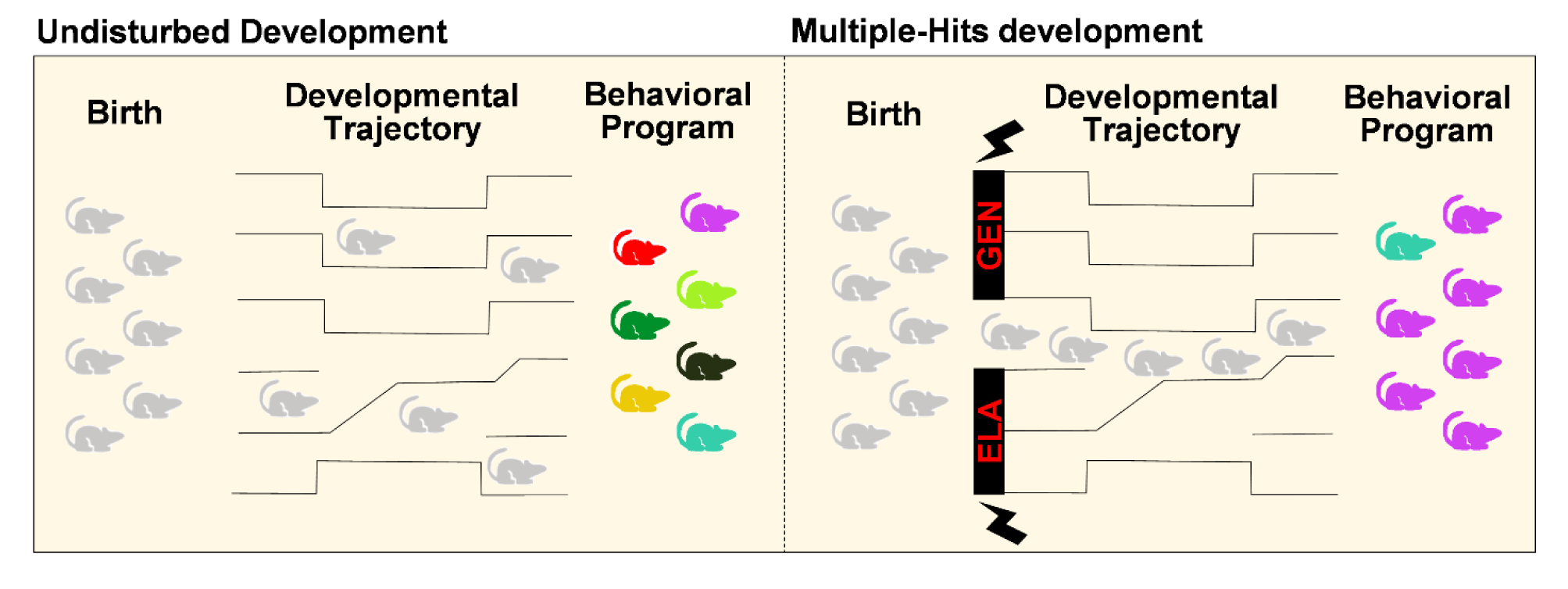
The combination of psychiatric vulnerabilities constrains behavioral complexity. Our results suggest that the convergence of multiple developmental hits limits the complexity of behavioral trajectories, constraining the individuals into stereotyped programs.

In addition, our work shows a previously undetected effect of *Cntnap2* haploinsufficiency, suggesting that early-life adverse experience can impact the behavioral outcomes based on the genetic predisposition. Further studies are needed to better understand what behavioral domains are affected by either the genetic component or the early-life experience.

One limitation of this study consists in the fact that the unbiased stratification of the dataset into two subpopulations does not match the original experimental groups. Moreover, neither of the two personalities perfectly corresponds to one of the experimental conditions, showing a maximum overlap of approximately 85%. This may be consequence of the limitations of measuring tools, flaws in the available statistical models, or undetected noise within the data. However, we believe that this percentage still provides enough evidence of the fundamental behavioral differences driven by genetic and environment vulnerabilities accounted in the study. Furthermore, we cannot exclude that the classification of *Cntnap2^+/-^*LBN mice within P1 may have some biological explanation. A minority of vulnerable subjects that express a behavioral profile similar to WT controls could be explained as statistically infrequent resiliency. As known, the adverse life experience resulting from scarcity of resources is mediated by maternal behavior (23, 25). Thus, it is reasonable to speculate that, despite the adverse environment, pups who received sufficient level of maternal care may escape the negative long-lasting effect of the mutation. On the other hand, mice raised in standard condition may not receive enough maternal care as the dam unevenly splits her attention across the whole litter, mimicking the effect of LBN and accounting for the ∼30% of subject in P2 that do not share the double-hit vulnerability.

Finally, we want to emphasize one aspect that falls at the intersection between the methodological approach and the philosophical conceptualization of preclinical research in mental illness. Among the many limitations of using animals for modeling complex psychiatric conditions, one critical aspect is the fact that the majority of behavioral tasks for rodents are highly context-dependent, forcing the animal to express specific traits when challenged with a non-conventional choice (3–5). However, the very nature of mental illnesses is to be chronic and context-independent (12, 13). Even in the case of stimulus-triggered phobia, or when the onset of a depressive phase is the result of a loss or trauma, the key defining point of the disorder is not the magnitude of the trigger, but the vulnerability of the patient to a specific emotional or behavioral state (12). With this work, we attempt to move beyond the limitations of a context-based approach by using a combination of standardized ethological screening and dimensional classification of complex phenotypic outcomes. This comes with multiple advantages: i) increased ecological-ethological validity, ii) Independency from pre-determined assumptions, iii) elimination of confirmation biases and iv) prioritization of qualitative evaluation, rather than quantitative oversimplification of the problem.

On the other hand, these methods partially lack interpretability of the obtained results. For instance, in our case, it is challenging to assign a satisfying ethological interpretation to different expressions of rearing behavior (BM2-7-8). A valuable solution would be to pair classical hypothesis-driven paradigm with data-driven methods to investigate as much as possible of the behavioral complexity.

Our results encourage the development of a new approach in which the long-term effects of genetic and environmental perinatal insults linked to neuropsychiatric disorders are analyzed with unbiased methodologies and with attention to individual profiles of behavioral subjectivity.

## METHODS AND MATERIALS

### Animals and housing

All mice were generated from our inbred *Cntnap2* colony with C57BL/6 background. Animals were housed in a 12h light/dark cycle with unrestricted access to food and water. A total of 38 age-matched adult (6 months old) mice (weight 25–35 g) were used in the study (n=20 males and 18 females, obtained from 3 litters raised in standard conditions and 4 litters raised in LBN). All efforts were made to minimize suffering of the animals. All experimental procedures were performed in accordance with Italian and European directives (DL 26/2014, EU 63/2010) and were reviewed and approved by the University of Trento animal care committee and Italian Ministry of Health (# 935/2021-PR).

#### Breeding strategy

To minimize the effect of confounding factors on the variables of interest (i.e. early-life stress and genotype), we adopted a standardize breeding strategy. Age-matched (postnatal age P=150-160 days) WT females were crossed with heterozygous males, eliminating the effect of maternal age and genotype on parental care. The first litter was discarded as to exclude the consequences of first experience in maternal care. Only second-born litters were included in the study.

#### Limited bedding and nesting

The LBN paradigm was adapted from Walker et al (23). At postnatal day 4 (P4) the dam with the entire litter was moved to an experimental cage. The cage featured an aluminum mesh placed at 2.5 cm distance from the base and nestled material reduced to ¼ with respect to a standard cage. The mother with the full litter was returned to a standard breeding cage once the pups reached P11. Control litters were moved from standard cages to standard cages at the same time-points, as to exclude the effect of experimental manipulation as a confounding factor. Both LBN mice and controls were weaned at postnatal day 21/22 and housed in standard cages with a single nestled square as an enrichment. Mice were then single-housed two weeks prior the beginning of the experimental pipeline.

### Behavioral Testing

#### Open Field

Each mouse was acclimated with the experimental room for 20 minutes. After habituation, each mouse was placed in the center of a large, squared arena (40 x 40 x 40 cm) and let free to navigate the open space for a total of 20 minutes. The experimental sessions were video recorded from above, using a camera interfaced with the Ethovision software (Noldus, Wageningen, the Netherlands) for the extraction of traditional behavioral descriptors. Two synchronized cameras in stereo-configuration were used for sidewise videorecording using the open-source software OBS studio and the resulting video analyzed using DeepLabCut 3D (33).

#### Classical assessment of the open field test

Distance traveled, velocity, time moving and time immobile were used as descriptors of locomotor activity. The ratio between the time spent in the center of the arena versus the borders (thigmotaxis) and the ratio between the time spent in the center of the arena versus the corners (safety preference) were used as a proxy for emotional state, as previously described.

#### Stimulus-evoked behavioral tracking in 3D for rodents (SEB3R)

SEB3R is a MATLAB pipeline for digitalized behavioral screening that was previously described in detail (2). The code and extensive instructions for SEB3R pipeline are available at the link: https://github.com/gchelini87/SEB3R. Briefly, 6 key-points along the mouse spine were tracked in DeepLabCut. Using the relative distances between key-points, SEB3R identifies a subset of statistically meaningful body-postures, for each animal. Then, similar postures are allocated within discrete behavioral categories that are shared across all (or most) subjects and labeled with a progressive enumeration (BMs). BMs labels are finally assigned to each frame of each video, matching the label of the original postures with the final label of the BMs.

### Statistical Analysis

The statistical analysis was computed using R-studio package in R software (*Integrated Development for R. RStudio, PBC, Boston, MA, USA*. http://www.rstudio.com/) or JMPpro17 software (SAS Institute Inc., Cary, NC, 2023). Graphical visualization was complemented by GraphPad Prism (GraphPad Software, Boston, MA, USA. www.graphpad.com) and Matlab 2021 (The MathWorks Inc., Natick, MA, USA. https://it.mathworks.com/products/matlab.html). Our statistical model excluded the effect of sex-differences; thus, we excluded the variable sex from the analysis. A two-way ANOVA was applied to test the main effect of *Genotype* (Gen), Early Life Experience (ELE) and their interactions for traditional descriptors and individual BMs. A two-way factorial ANOVA with multiple response variables (all BMs) was used to investigate the overall effect of the dependent variables on the entire behavioral structure. Post-hoc multiple comparisons were calculated using Tuckey’s corrections. Multivariate analysis (MANOVA) was used to evaluate BMs co-variation across experimental groups. While no significant impact of independent variables was detected, the resulting canonical scores highlighted a clear misalignment of the experimental groups as shown in Fig. 2. The suitability of the data for multivariate testing was confirmed by inspecting the QQ plots of residuals.

#### K-mean clustering

To stratify the study cohort into distinct behavioral profiles we ran an iterative *k-clustering* algorithm testing from 2 to n (=38) possible combination of data repartition. The cubic clustering criterion was used to determine the best ft to the data and the classification reliability was confirmed using a two-way ANOVA of the distance from centroids.

#### Network Analysis

Network structure and analysis of centrality was performed using the igraph library package (https://github.com/igraph/igraphdata) contained in R-studio software. For each behavioral profile (P1 and P2), we calculated the pairwise Spearman correlation (*ρ*) of BMs expression between each subject, obtaining a squared correlation matrix (Fig. 4A). The raw correlation values were then transformed into binary entries (0 or 1) using a value of *ρ=0.8* as a cutoff; every correlation above 0.8 was converted to 1, while the others were set to 0, generating an adjacency matrix. Next, undirected networks were designed starting from the adjacency matrices. Each node within the networks represents a single mouse and the connecting line the degree of similarity between connected subjects. Then, the following centrality measures were extracted for each subject: barycenter centrality, betweenness, clustering coefficient and mean path distance. Using the “proper centralities” function from the R-studio pipeline Central Informative Nodes in Network Analysis (CINNA), we selected the most informative centrality indicators identified after dimensional reduction (https://cran.r-project.org/web/packages/CINNA/index.html).

## Acknowledgments

We thank all the administrative and technical staff of CIMeC for support. A special thank goes to Mrs. Michela Maffei, animal caretaker of the CIMeC animal facility, for her endless care and support provided in managing the animal colony used in the study and Dr. Tommaso Pecchia for helping with the technical implementations of some of the behavioral apparatuses. GC effort was covered by the ‘CARITRO postdoctoral fellowship’, funded by Fondazione Cassa di Risparmio di Trento e Rovereto. AS acknowledges funding from Next Generation EU, in the context of the National Recovery and Resilience Plan, Investment PE8 – Project Age-It: “Ageing Well in an Ageing Society” [DM 1557 11.10.2022]. This resource was co-financed by the Next Generation EU.

## Author Contributions

V.CA. performed the data analysis and wrote the first version of the manuscript. V.CI. contributed to research work. N.B. contributed to the conceptualization of the work. A.S. and Y.B. contributed to the conceptualization of the work, provided funding, and edited the manuscript. G.C. conceptualized and supervised the work, performed all the experiment, provided funding, and wrote the manuscript.

## Supplementary Figures

**Supplementary figure 1.**
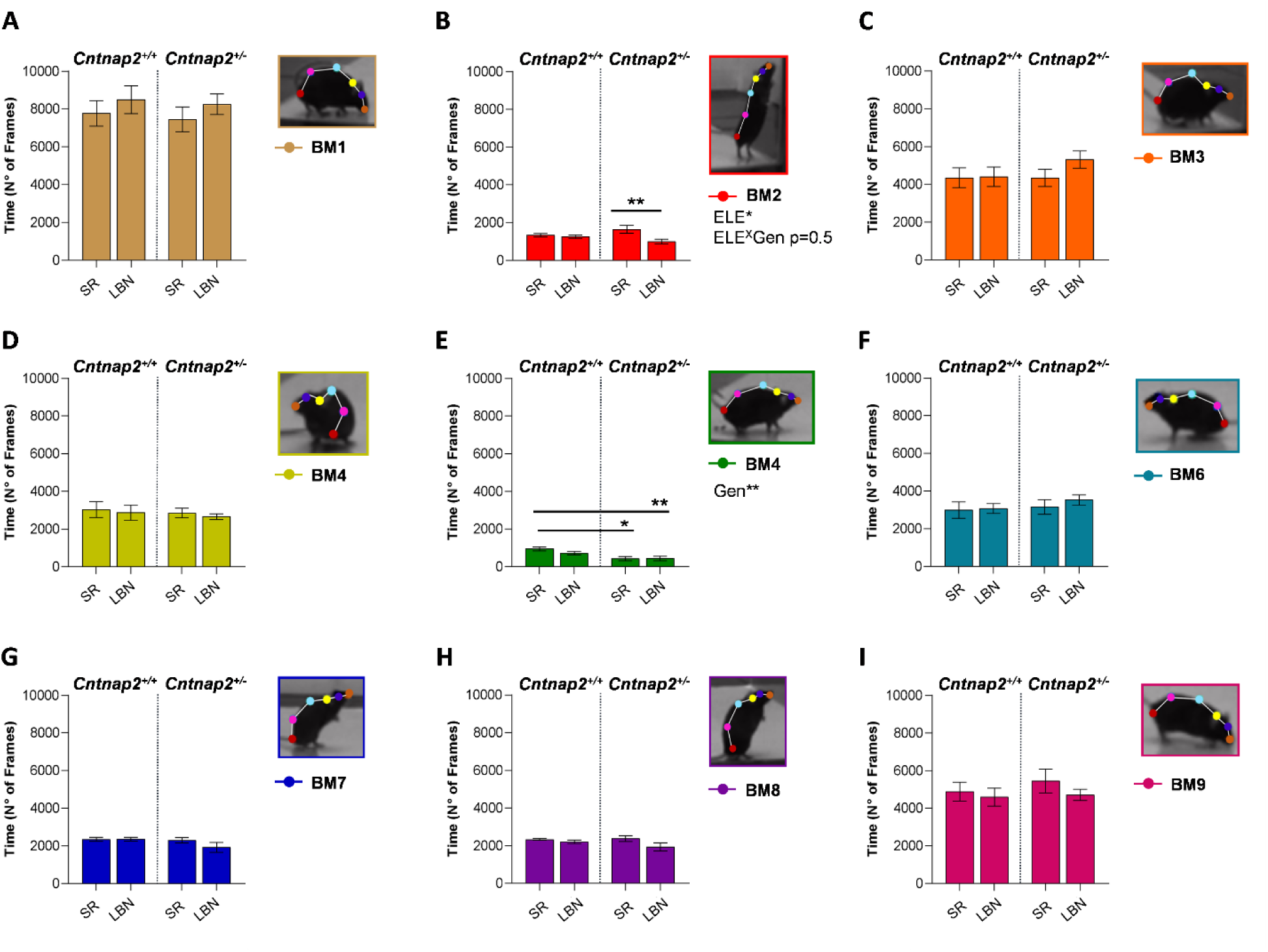
Moderate variation in single BMs expression observed across experimental groups. **A-I)** The plots report the BMs expression in number of frames. **B)** Marginal effect of the ELE was found in BM2 (p=0. 014,) with a near to significance effect of the ELE^X^Gen interaction (p=0.051). The effect was mostly driven by significant differences between *Cntnap2*^+/-^ SR and LBN (p=0.009). **E)** A prominent effect of the genotype was also discovered in BM5 (p=0.001), showing significant reduction in both Cntnap2^+/-^ groups compared to *Cntnap2*^+/+^SR (*Cntnap2*^+/+^SR vs *Cntnap2*^+/-^SR p=0.02, vs *Cntnap2*^+/-^LBN p=0.009). Data are expressed as mean +/- SEM. Statistical analysis was performed by two-way ANOVA. Post-hoc analysis was carried out using Tuckey’s correction for multiple comparisons. *p<0.05 / **p<0.001.

**Supplementary Figure 2.**
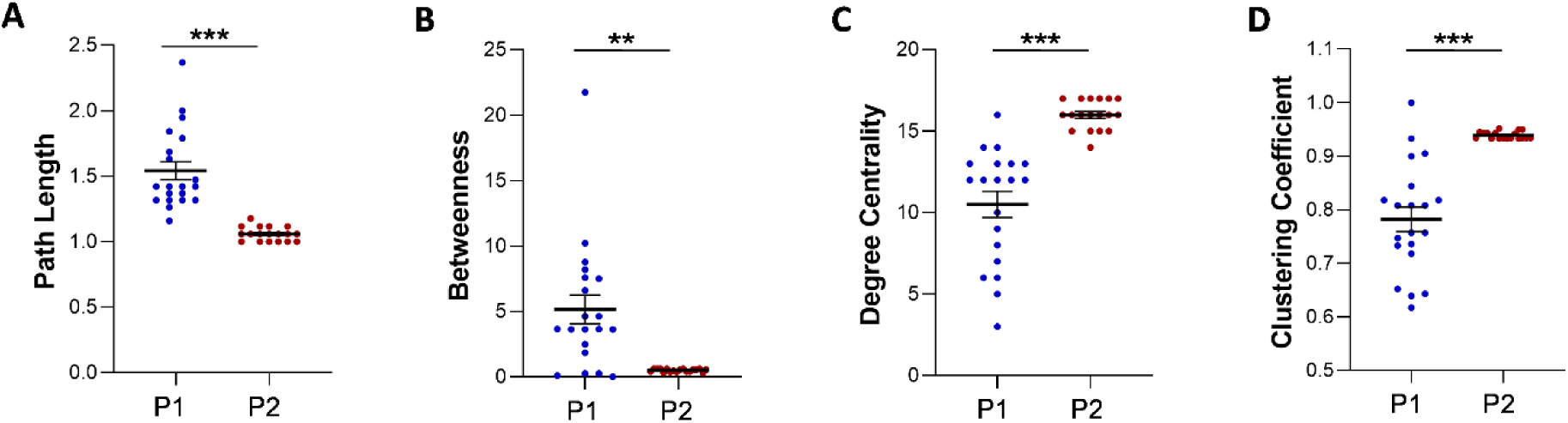
Additional centrality measure showing marked differences in the network homogeneity. Using multiple parameters: **(A)** Path length, **(B)** Betweenness, **(C)** Degree of centrality and **(D)** Clustering Coefficient, we compared the homogeneity of P1 and P2, obtaining identical results that indicate higher similarity among P2 subjects. Mann-Whitney test. **p<0.001 / ***p<0.0001.

